# Multi-corneal barriers-on-a-chip to recapitulate eye blinking shear stress forces

**DOI:** 10.1101/2019.12.23.886242

**Authors:** Rodi Abdalkader, Ken-ichiro Kamei

## Abstract

Human corneal epithelium coexists tear fluids and shows its barrier functionality under the dynamic conditions of eye blinking. However, the current *in vitro* cell culture settings for corneal epithelial cells lack the dynamic flow conditions to recapitulate shear stress of eye blinking, hindering corneal function evaluation. We developed a microfluidic platform enabling the dynamic culture of the human corneal barrier with recapitulation of eye blinking. The device consisted of upper and lower channels separated by a porous membrane. Human corneal epithelial cells (HCE-T) were seeded on a porous membrane in an upper channel and cultured for ten days. The cells formed a barrier with high expression of zonula occludens 1 (ZO-1) tight junction protein on day seven, and the translocation of fluorescein sodium across the barrier in the microfluidic device was comparable to that in the transwell system. Then, bidirectional and unidirectional flows were applied in the upper and lower channels, respectively, and the cells in the upper channels were stimulated with 0.6 dyn s cm^-2^ of shear stress. While the fluid stimuli after 24 h did not affect cell adhesion, the flow stimuli facilitated the expression of cytokeratin 19 (CK-19) intermediate filaments in cells after 24 h, indicating strengthening of the barrier function. Furthermore, the morphological single-cell analysis revealed an increase in cell body area rather than nuclei. We envision that this multi-corneal barriers-on-a-chip device will unlock new possibilities in ophthalmic drug development and will be useful for studying the mechanobiology of the ocular surface.

## Introduction

The human cornea is the outer most refractive layer of the eye. It is composed of layers of epithelial cells supported by a collagen-rich extracellular matrix that contains keratocytes.^1^ The corneal epithelium is usually the main barrier for ophthalmic drug permeability and toxicity studies.^2^ These studies have been conducted *in vitro* with 2D models using established corneal cell lines or *in vivo*, particularly using rabbits.^3^ *In vitro* models lack the 3D structure of the cornea, which can affect the evaluation of the biological function of the barrier.^4^ Moreover, they are performed under static conditions that are different from the *in vivo* dynamic conditions during eye blinking. Blinking dynamic forces have two crucial aspects. The first one is related to drug toxicity and distribution. In this case, the rabbit eye model can mislead the interpretation of pre-clinical test results because, compared to humans, the low blinking rate in rabbits can cause higher interaction between the drug and the ocular surface, leading to higher drug permeation.^5,6^ The second is related to the corneal surface homeostasis and functions under the shear stress stimuli during eye blinking.^7^ To conduct proper investigation of the shear stress effects, we need a platform that is structurally relevant to the human cornea, i.e., one that contains an apical side that holds the tear fluid of the eyes and a proximal side that interconnects with the aqueous humor. Thus, the ideal platform should have the ability to recapitulate the ocular fluid dynamic flow and shear stress forces occurring during eye blinking. This system should additionally allow multi-tests in terms of side-by-side comparisons within the model, which can be used for further mechanobiological studies and drug screening experiments.

Recently, the organ-on-a-chip technology has become an attractive solution for the recapitulation of the complexity of a variety of organs such as gut^8^, liver^9^, and lung^10^. Multi-layers of cells can be grown in microfluidic devices in a 3D manner supported by sufficient niches of extracellular matrices.^11,12^ Microfluidic systems can also provide a unique solution for mimicking the fluids dynamics in organs under controlled parameters of shear stress and fluids velocity.^13,14^ Microfluidic devices have been used to study the mechanobiology of organs *in vitro,* in particular the influence of blood flow-mediated shear stress on endothelial cell deformation and morphology in healthy and diseased situations.^15,16^ However, the reported microfluidic platforms are not sufficiently developed for evaluating the effect of shear stress forces on the homeostasis of the corneal epithelial barrier during eye blinking.

Here, we developed a multi-corneal barriers-on-a-chip for growing human corneal epithelial cells under the dynamic flow stimuli that can recapitulate the shear stress forces during eye blinking (Fig. 1). We studied the creation and development of biological features of the corneal epithelial barrier in the microfluidic device in comparison with the conventional transwell system. To mimic the eye blinking *in vitro*, we have applied bidirectional flow on the apical side of the cells and a continuous flow on the proximal side for 12–24 hours. Then, we investigated the influence of flow dynamics on human corneal cell functions, phenotypes, and morphological changes.

**Fig 1.**
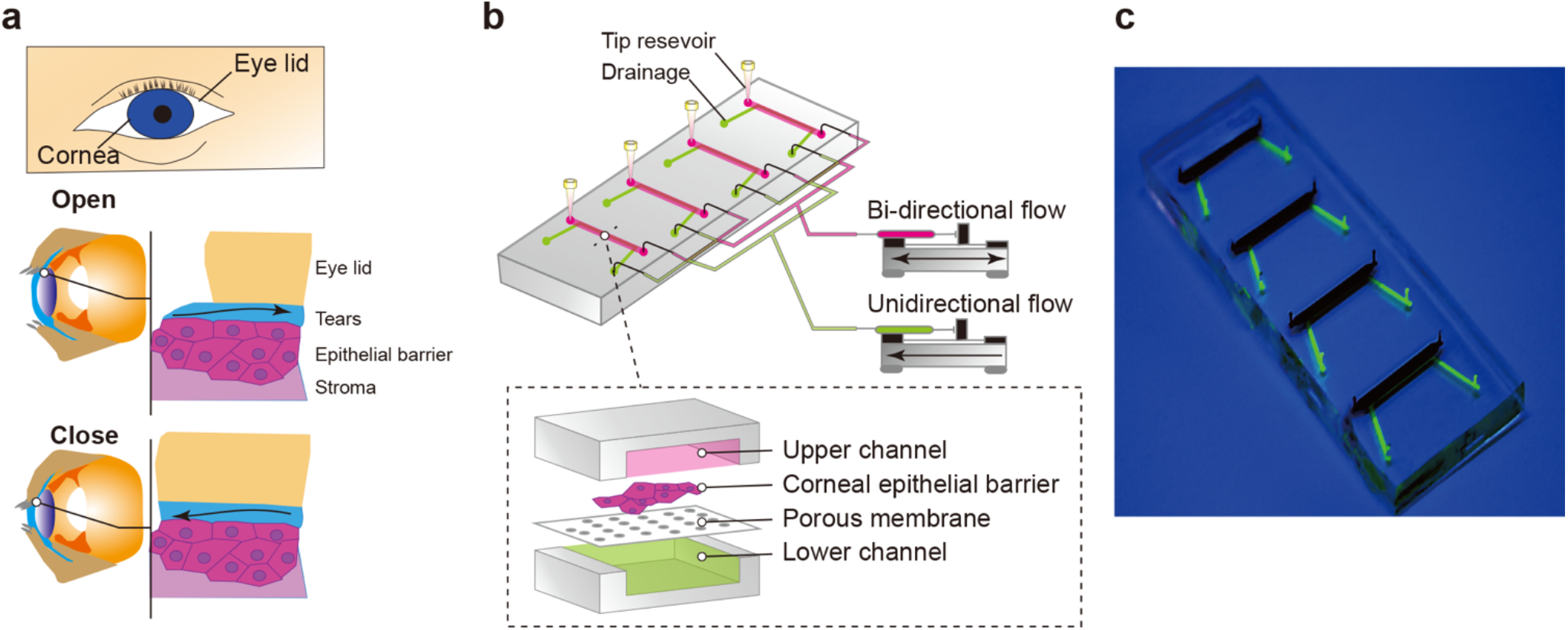
Conceptual illustration of the multi-corneal barriers-on-a-chip. **a**, Illustration of the human corneal structure showing the dynamic of tear film movement during eye blinking. **b**, Schematic diagram of the dynamic flow perfusion system. The upper channels were connected with a syringe pump that generates a bidirectional flow, while the lower channels were connected to another syringe pump that generates a unidirectional flow. Microvolume pipet tips were fixed in the outlets of the upper chamber to serve as a reservoir. **c**, The final structure of the microfluidic device.

## Experimental

### 1. Microfluidic device fabrication and characteristics

The microfluidic device on-a-chip was fabricated using stereolithographic 3D-printing techniques and solution cast-molding processes.^17^ The mold for the microfluidic channels was produced using a 3D printer (Keyence Corporation, Osaka, Japan). Two molds were fabricated: the upper block and the lower block. Each block contained four chambers (15 mm length, 1.5 mm width, and 0.5 mm height). Prior to use, the surfaces of the molds were coated with trichloro(1H,1H,2H,2H-perfluorooctyl) silane (Sigma-Aldrich, St. Louis, MO, USA). Sylgard 184 PDMS two-part elastomer (ratio of 10:1 pre-polymer to curing agent; Dow Corning Corporation, Midland, MI, USA) was mixed, poured into the molds to produce a 4-mm-thick PDMS upper layer and a 0.5-mm-thick PDMS lower layer, and degassed by using a vacuum desiccator for 1 h. The PDMS of the lower block was fixed on a glass slide. The PDMS material was then cured in an oven at 65°C for 24 h. After curing, the PDMS form was removed from the molds, trimmed, and cleaned. A clear polyester (PET) membrane (pore size of 0.4 μm, thickness of 10 μm, and nominal pore density of 4 × 10^6^ pores cm^-2^) was fixed on each chamber of the lower PDMS block. Both PDMS blocks were corona-plasma-treated (Kasuga Denki, Inc., Kawasaki, Japan) and bonded together by baking in an oven at 80°C. Scanning electron micrographs were obtained using a JCM-5000 microscope (JEOL Ltd., Tokyo, Japan) using 10 kV. Prior to imaging, a 5-nm-thick platinum layer was sputtered on the samples (MSP 30 T; Shinku Device, Sagamihara, Japan).

### 2. Fluid diffusion and translocation in the microfluidic device

Fluorescein sodium and rhodamine B in a concentration of 250 μg mL^-1^ were dissolved separately in Hanks balanced buffer supplemented with calcium (HBSS^+^). The rhodamine B solution was added to the upper chamber, while the lower chamber was filed fluorescein sodium. The translocation between both fluorescent dyes was recorded for 15 min using fluorescence microscopy (Nikon, Tokyo, Japan).

### 3. Human corneal epithelial cell culture

HCE-T cells were provided by RIKEN Bio-resource Research Centre (Ibraki, Japan). Cells were cultured with DMEM/F12 supplemented with 5% (v/v) fetal bovine serum (FBS), 5 μg mL^-1^ insulin, 10 ng mL^-1^ human epithelial growth factor (hEGF), and 0.5% dimethyl sulfoxide (DMSO). The cells were passaged with trypsin-EDTA (0.25–0.02%) solutions at a 1:2 to 1:4 subculture ratio.

### 4. Human corneal epithelial barrier construction in the microfluidic device

Prior to use, a microfluidic cell culture device was placed under ultraviolet light in a biosafety cabinet for 30 min. The microfluidic channels were washed with DMEM/HamF12. Cells were harvested using trypsin and collected in a 15-mL tube. Following centrifugation, the cells were resuspended in DMEM/HamF12 medium and introduced into the upper channel of the microfluidic device via a cell inlet with a cross-sectional area of 0.23 cm^2^ at a density of 1 × 10^6^ cells mL^-1^. The microfluidic devices were placed in a humidified incubator at 37°C with 5% CO_2_ for 5, 7, and 10 days. The medium in each chamber was periodically changed every 24 h. For the transwell system, 1 × 10^5^ cells in 0.5 mL of medium were added to the 24-mm transwell (0.4 mm pore, 1.16 cm^2^ area, Corning). The lower receiver channel was filled with 1.5 mL of DMEM/HamF12. The cells were placed in a humidified incubator at 37°C with 5% CO_2_ and the medium was changed every two days.

### 5. Barrier morphology and functions

#### 5.1. Immunocytochemistry and microscopic imaging

Cells were fixed with 4% paraformaldehyde in PBS for 25 min at 25°C and then permeabilized with 0.5% Triton X-100 in PBS for 10 min at 25°C. Subsequently, cells were blocked with blocking buffer (5% (v/v) normal goat serum, 5% (v/v) normal donkey serum, 3% (w/v) bovine serum albumin, 0.1% (v/v) Tween-20) at 4°C for 24 h and then incubated at 4°C for overnight with the primary antibody (zonula occludens 1 (ZO-1) rabbit anti-human or cytokeratin 19 (CK19) mouse anti-human) in blocking buffer. Cells were then incubated at 37°C for 60 min with a secondary antibody (AlexaFluor 488 donkey anti-rabbit IgG or AlexaFluor 599 donkey anti-mouse IgG, 1:1000; Jackson Immuno Research, West Grove, PA, USA) in blocking buffer prior to a final incubation with 4’,6-diamidino-2-phenylindole (DAPI) at 25°C. For imaging, we used a Nikon ECLIPSE Ti inverted fluorescence microscope equipped with a CFI plan fluor 10×/0.30 N.A. objective lens (Nikon, Tokyo, Japan), CCD camera (ORCA-R2; Hamamatsu Photonics, Hamamatsu City, Japan), mercury lamp (Intensilight; Nikon), XYZ automated stage (Ti-S-ER motorized stage with encoders; Nikon), and filter cubes for fluorescence channels (DAPI, GFP and CY5; Nikon). The barrier thickness was further investigated by the confocal laser scanning microscopy (A1R, Nikon). The z scanning mode was applied from top to bottom. The z images were then analyzed using ImageJ software (National institute of health, Maryland, USA).

#### 5.2. Fluorescein sodium permeability

The permeability experiments were performed in HBSS^+^ (pH 7). Test samples were prepared by diluting the fluorescein sodium solution in HBSS^+^ to a concentration of 250 μg mL^-1^. In the transwell system, prior to the experiment, the cells were washed with HBSS and incubated with 400 μL and 2000 μL of HBSS^+^ in the donor and receptor compartments, respectively, during the subsequent 30 min. At the start of the experiment, HBSS^+^ was removed from the donor compartment, and 400 μL of fluorescein sodium was added. Samples (200 μL) were taken from the receptor compartment at regular time intervals over 120 min and replaced with the same volume of fresh buffer.

In the microfluidic device, the permeability assay was conducted by perfusion of both the upper and lower channels with culturing medium at a flow rate of 0.01 mL min^-1^. The lower channels were perfused with HBSS^+^ and the upper channels were perfused with fluorescein sodium solution. Fluorescence (485 nm excitation and 530 nm emission) was detected on a plate reader (Synergy HTX, Bio Tek Instruments, Winooski, USA). The steady-state flux (J) values across the corneal barrier were determined from the linear ascents of the permeation graphs by means of the relationship:

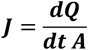

The permeation coefficient, P_app_, was calculated as:

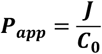

where C_0_ represents the initial fluorescein sodium concentration in the donor compartment, Q is the quantity of substance crossing the corneal epithelial barrier, A is the exposed corneal area, and t is the exposure time.

### 6. Dynamic flow stimuli

The upper channels were exposed to a bidirectional flow using a syringe pump apparatus (KD Scientific, Fisher scientific, Holliston, USA), and the dynamic flow model consisted of three phases. Phase 1: a withdraw flow at a rate of 5 μL s^-1^ for 20 s. Phase 2: an infuse flow at a rate of 5 μL s^-1^ for 20 s. Phase 3: an infuse flow at a rate of 0.1 μL min^-1^ for 120 s. The lower channels were connected to another syringe pump apparatus with a continuous unidirectional flow of 0.1 μL min^-1^. The shear stress in the microchannels was calculated using the equation:

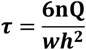

where n is the viscosity of DMEM at 37°C (8.91 × 10^-3^ dyn s cm^-2^), Q is the flow rate in cm^3^ s-^1^, w is the width of the microfluidic channel in cm, and h is the height of the microfluidic channel in cm.

Single-cell profiling based on fluorescent microscopic images was conducted using the CellProfiler software (Version 3.1.8; Broad Institute of Harvard and MIT, USA^18^). The Otsu’s method was employed to identify cells, and the fluorescence signal from individual whole cells, cytoplasm, and nuclei were separately quantified automatically. Further analysis of single-cell morphological descriptors was performed by t-Distributed Stochastic Neighbour Embedding (t-SNE) techniques using the open-source Orange 3 software (Version 3.23.1; Bioinformatics Laboratory, Faculty of Computer and Information Science, University of Ljubljana, Slovenia^19^). Cell morphology was evaluated by determining the aspect ratio (AR) through the equation:

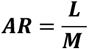

where L is the cell major axis length and M is the minor axis length.

### 7. Statistical analysis

All experiment represented as mean ± S.D. The unpaired t test, nested t test, or Tukeys’s multiple comparison test were performed using GraphPad prism 8 (GraphPad Software, La Jolla California, USA)

## Results and discussion

### Fabrication and characteristics of the multi-well microfluidic device

The design of the multi-well microfluidic device is shown in Fig. 1. The device has four upper and lower channels separated by a clear polyester (PET) porous membrane. We have developed a simple yet uniquely practical process for the device fabrication (Supplementary Fig. 1). To ease the removal of PDMS blocks from the molds, trichloro silane was utilized. The devices are finally sealed by bonding without inducing damage to the porous membrane. SEM images of the device showed the intactness of the porous membrane confined between the upper and lower PDMS blocks (Supplementary Fig. 2). To test liquid leakage from the microchannels, commercially available colored dyes were used. Visual observations confirmed no leakage from the microchannels for 150 min, indicating that the porous membranes were completely sealed in between the upper and lower channels; no side leakage of the dyes was noticed (Supplementary Fig. 3).

To evaluate the liquid translocations between the upper and lower channels, the hydrophilic fluorescent dye (fluorescein sodium, rhodamine B) translocation was tested for 15 min (Fig. 2a). The rhodamine B fluorescence signal was clearly observed after 5 min and continued to increase in time-dependent manner. On the other hand, the fluorescein sodium fluorescence signal was gradually reduced. These results indicate the active diffusion of liquids between the upper channels and the lower channels and vice versa. (Fig. 2b).

**Fig. 2.**
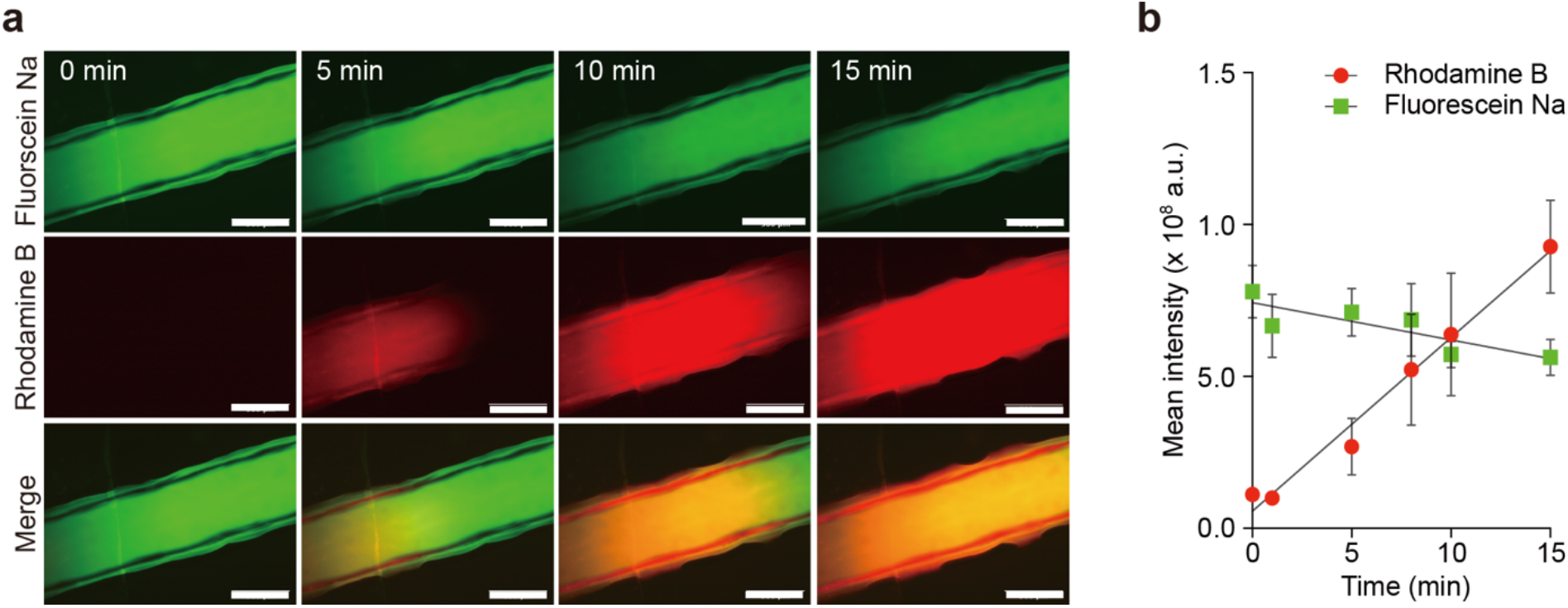
Fluids translocation in the multi-well microfluidic device. **a**,Representative fluorescence images of rhodamine B (red) and fluorescein sodium translocation in the microfluidic channels (green) during 15 min. The rhodamine B solution was added into the upper channels, while fluorescein sodium was added into the lower channels. Scale bar, 500 μm. **b**, Increase of fluorescence intensity of rhodamine B in the lower channels plotted against the decrease of fluorescein sodium. Data are represented as means ± SD of three independent experiments.

### Culturing of corneal epithelial cells and their ability to form a barrier

In order to evaluate the growth and formation of the corneal epithelial barrier, HCE-T cells were seeded in the microfluidic devices at a density of 6 × 10^3^ cells cm^-2^, which is lower than the density of cells used in the conventional transwell system (0.89 × 10^5^ cells cm^-2^) (Supplementary Fig. 5). The barrier was formed starting from day 5. On day 7, the microfluidic channels were fully covered with cells. Clusters of cells were visible, forming a second layer of cells. The ZO-1 expression was clearly increased on day 7 and then decreased on day 10 (Fig. 3a and b). Moreover, cells grown in the microfluidic device formed a thicker barrier compared to those grown in the transwell system (Fig. 3c).

**Fig. 3.**
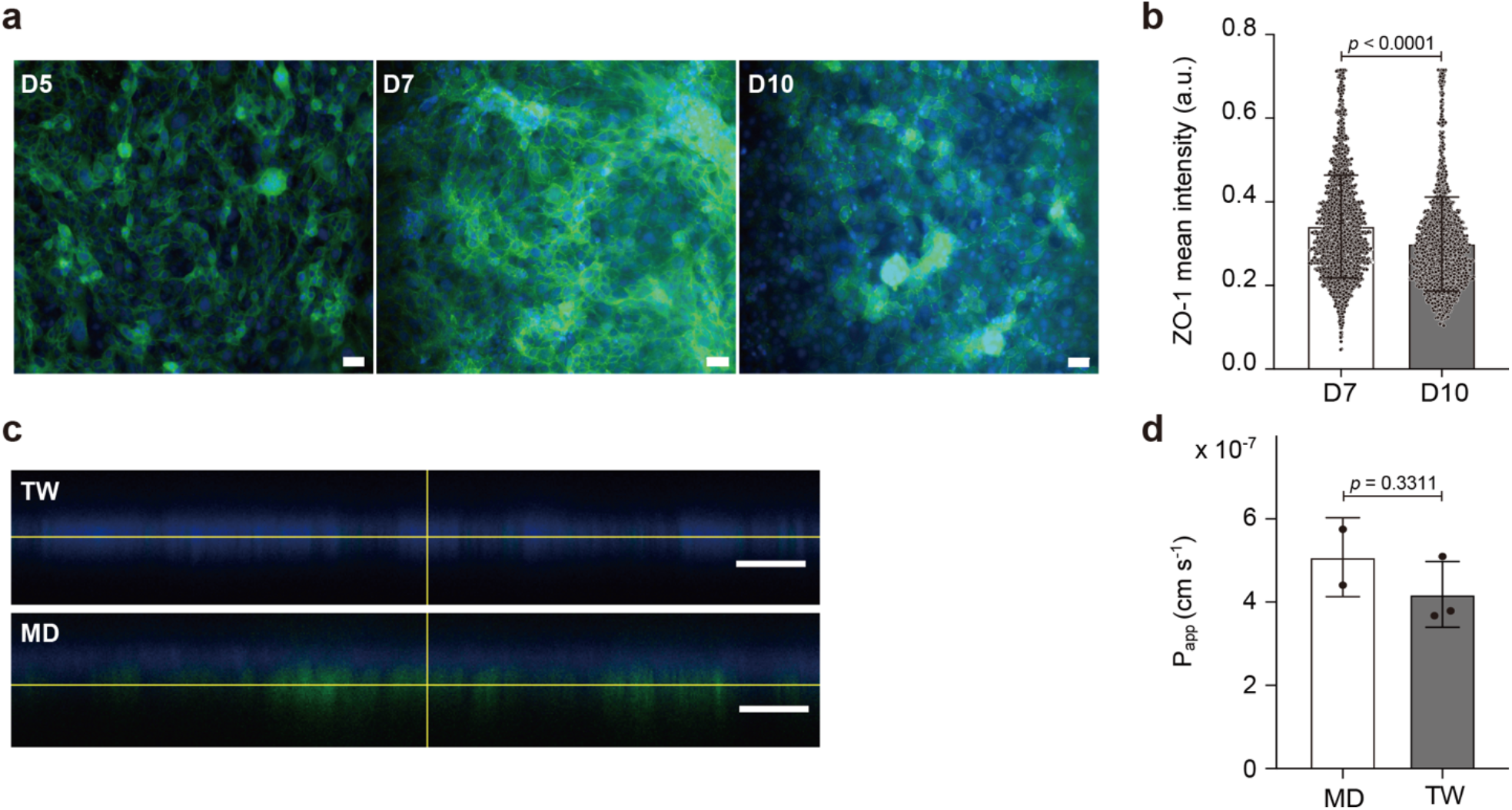
Characterization of the corneal epithelial barrier in the multi-corneal barriers on-a-chip. **a**, Immunofluorescence staining of ZO-1 in HCE-T human corneal epithelial cells grown in the multi-well microfluidic devices at days 5, 7, and 10. Green, ZO-1; Blue, DAPI. Scale bar, 50 μm. **b**, Quantitative single cell profiling of ZO-1 expression in HCE-T cultured in the multi-well microfluidic device at days 7 and 10. Fluorescent intensity was obtained from images of three independent experiments; 1800 cells were randomly selected and analyzed for each sample. **c**, Z-stack images of the corneal epithelial barrier showing ZO-1 expression in the transwell (TW) at day 7 and in the multi-well microfluidic device (MD) at day 5. Green, ZO-1; Blue, DAPI. Scale bar, 100 μm. **d**, Fluorescein sodium permeability in the corneal epithelial barrier grown for 7 days in the multi-well microfluidic device. Fluorescein sodium solution was added to the upper channels, while the lower channels were filled with HBSS buffer. A flow of 0.01 mL min^-1^ was applied. Samples were collected from the lower chambers. Barrier permeability is presented as apparent permeability (Papp). Data are represented as mean ± SD. The *p*-values were determined by the unpaired *t* test.

To confirm the integrity of the corneal epithelial barrier in the microfluidic device, we tested the permeability rate of fluorescein sodium after 7 days of cell culturing. The P_app_ values of fluorescein sodium in the microfluidic device and transwell system were 5.07 × 10^-7^ ± 9.4 × 10^-8^ and 4.18 × 10^-7^ ± 7.9 × 10^-8^, respectively (Fig. 3d). There was no significant difference in the permeability of fluorescein sodium between the barriers in the microfluidic devices and that in the transwell system, indicating that the epithelial barrier intactness in the microfluidic device is equivalent to that in the transwell system (*p*-value **=** 0.3311).

HCE-T cells grown in the transwell system can form an intact barrier between day 7 and day 15 with transepithelial electric resistance (TEER) values ranging 400–1000 Ω cm^-2^.^20^ Our result show that the TEER of HCE-T cells in the transwell system gradually increased after seeding and reached 499.52 ± 22 Ω cm^-2^ after 7 days (Supplementary Fig. 4). In the current study, the TEER measurement in the multi-well microfluidic device was not conducted but instead the barrier intactness was confirmed by measuring the fluorescein sodium permeability. The correlation between the permeability of fluorescein sodium and TEER measurement has been previously reported in a Caco-2 cell barrier.^21^ Moreover, although there was a significant increase in the ZO-1 staining that led to the increase of the barrier thickness in the microfluidic channels, the fluorescein sodium permeability was not reduced compared to that in the transwell system. This can be due to the active transport of fluorescein sodium in both cellular and paracellular pathways.^22^ In addition, the incomplete multi-layer formation will not influence the fluorescein sodium permeability. Regarding the corneal barrier thickness in the microfluidic device, cells formed clusters that led to almost two layers of cells after seven days of culturing. In contrast, only a monolayer of cells was found in the transwell system. HCE-T cells tend to form 2–3 layers of cells when left in an air-liquid interface.^23^ In our study, the air-liquid interface was avoided to limit the changes in cell phenotype and morphology to the effect of flow dynamic stimuli.

### The effect of fluid dynamic flow stimuli on cellular adhesion, functions, morphology, and phenotype

One eye blinking cycle consists of withdraw and infuse flow for a duration of 40 s. Thus, to create the fluid dynamic stimuli in the microfluidic device, two different types of flow were applied: the bidirectional flow consisted of withdraw-infuse circles applied to the upper channels, and the unidirectional infuse flow was only applied to the lower channels. The bidirectional flow induced a shear stress of 0.6 dyn s cm^-2^ in the microfluidic channel.

In order to investigate the effect of shear stress mediated by fluid dynamics on corneal cell adherence, the cell nuclei were counted after 12 and 24 h. No significant reduction of nucleus counts was observed in neither of the conditions (dynamic or static) (Fig. 4).

**Fig. 4.**
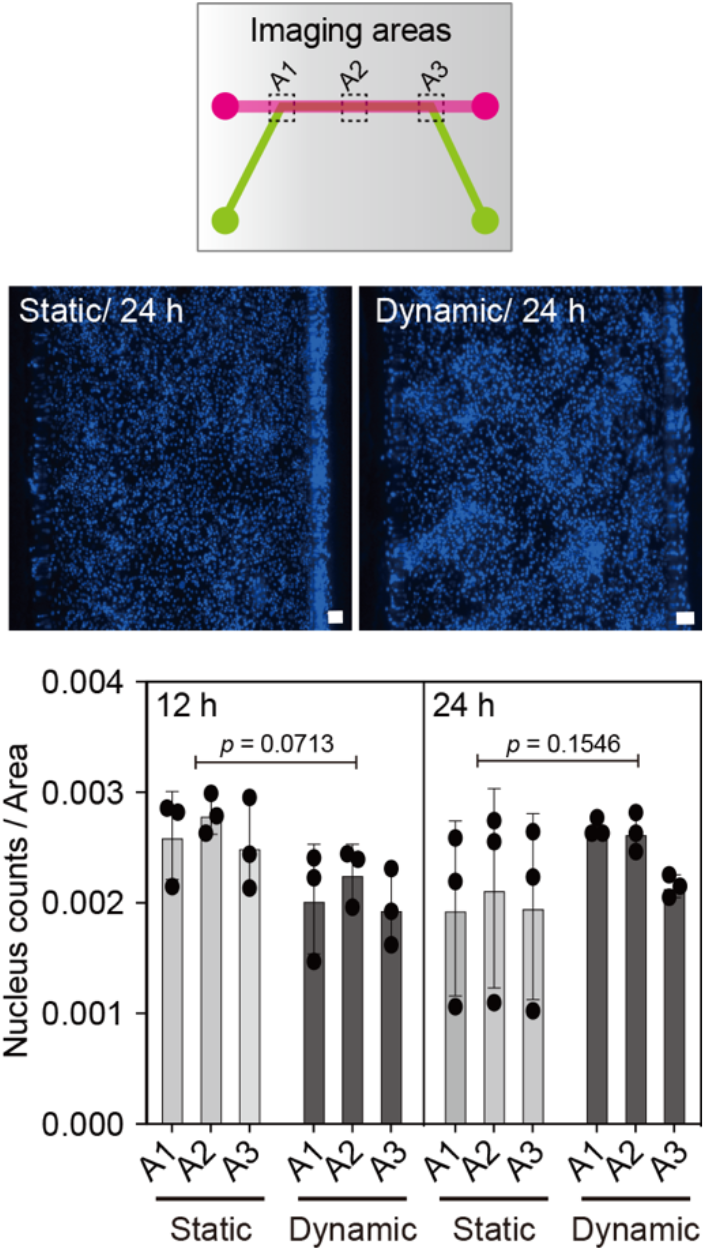
4 Dynamic flow stimuli in the multi-well microfluidic device. HCE-T cell adherence under dynamic flow stimuli. After 12 and 24 h, the cells were fixed and stained with the nucleus stain DAPI (blue). Scale bar, 100 μm. Images from three different areas of the microchannels were obtained: near to the inlets, in the middle, and near to the outlets. Nucleus count was performed and the total nucleus number was normalized by the image acquisition area. Data are represented as mean ± SD. The *p*-values were determined by the Tukey test.

To test the effect of the fluid dynamic stimuli in the microfluidic device on cell functions, the expression of CK-19 was investigated. At 24 h after the application of dynamic flow stimuli, the expression of CK-19 was significantly increased compared to the cells kept in static conditions (*p*-value <0.0001) (Fig. 5a and c). Previously, it was found that the application of a shear stress of 4 dyn s cm^-2^ on a monolayer of human corneal epithelial cells increased the amount of actin filaments in cells and that it has shortened the wound healing time.^24^ Actin filaments are known for their mechanoresponsive properties in different cells. Together with myosin, actin provides the contractile forces that initiate cell migration.^25^ Cytokeratins, on the other hand, are intermediate filaments that maintain the flexibility and elasticity of cells.^26^ They are important for keeping the integrity of the corneal barriers. Disfunction in keratins causes serious ocular surface deficiencies.^26^ Our result suggests that the application of dynamic flow stimuli on the corneal barrier could reorganize and boost CK-19 to counteract the shear stress forces.

**Fig. 5.**
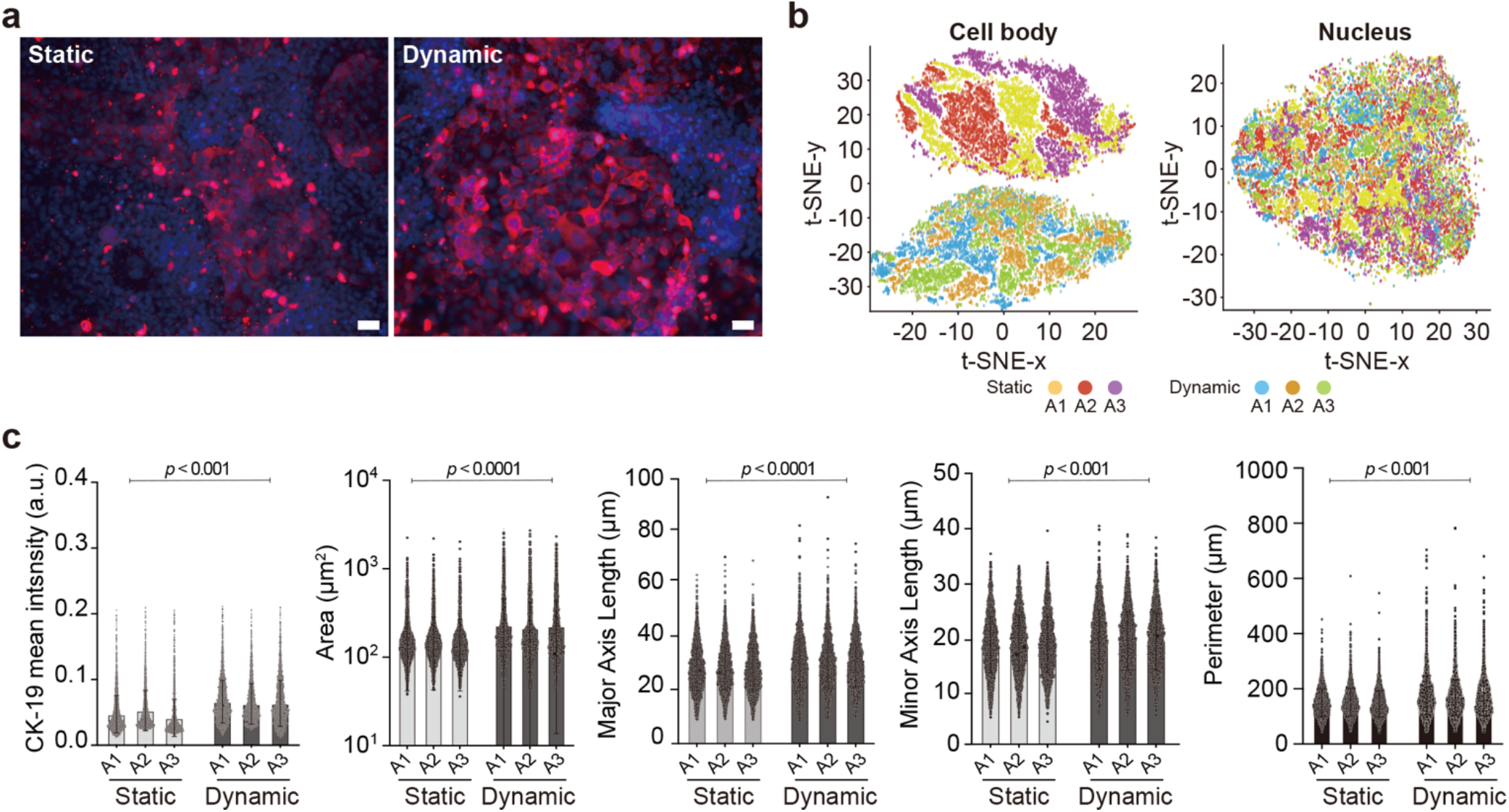
The effect of the dynamic flow stimuli on cell functions and morphology. **a**, Immunofluorescence staining of CK-19 in HCE-T human corneal epithelial cells exposed to dynamic flow stimuli for 24 h. Pink, CK-19; Blue, DAPI. Scale bar, 50 μm. **b**, t-SNE analysis of HCE-T cell populations. Over 100 cell features (morphology, intensity, and texture) were extracted from images of three independent experiments. **c**, Quantitative single-cell profiling of CK-19 expression in HCE-T cells and the related morphological features under static and dynamic flow conditions. Analysis was performed on images of three independent experiments; 3000 cells were randomly selected and analyzed for each sample. Data are represented as mean ± SD. The *p*-values (<0.001, 0.0001) were determined by the nested t test.

To further evaluate the morphological changes of cells under dynamic conditions, single-cell profiling was conducted. The assessed features included the morphological descriptors of the nucleus and cell body in addition to cell surface texture and the intensities of CK-19 and DAPI staining. t-SNE analysis showed a clear separation between the cell body populations of cells exposed to dynamic flow stimuli and those kept at static conditions. Moreover, cells in different areas of the microfluidic channels showed homogenous phenotype change. On the other hand, t-SNE analysis did not show any clear separation among the populations of nuclei under dynamic flow stimuli or under static conditions (Fig. 5b). This result indicates that shear stress mediated by dynamic flow stimuli in the microfluidic device has led to a morphological alteration in cell body rather than in the nucleus. The aspect ratio of the cells under dynamic flow stimuli was significantly increased compared to cells under static conditions (*p*-value < 0.001) (Supplementary Table. 1). This result confirms that the cell body has been stretched following application of dynamic flow stimuli. Histological analysis of human cornea structure shows a distinct difference between the morphology of superficial epithelial cells, wing cells, and the lower basal cells,^27^ although there might be a development-based reason for these variations. Our results provide an evidence for the role of shear stress during the application of a dynamic flow, and its underlying mechano-signaling mechanism may have induced changes in the morphology of cells, which became more flat in a way similar to the human corneal superficial cells.

These investigations show the necessity of blinking stimuli in the cornea biomimetic model for mimicking of the mechanosensory trigger needed for the recapitulation of the corneal barrier. A limitation of our study includes the short application dynamic flow stimuli (24 h). Further studies must consider the application of blinking stimuli for longer periods of time. Additionally, collective changes in the corneal cells in different layers of the barrier should be evaluated. Despite these limitations, the multi-well microfluidic device provides a valuable tool for the recapitulation of the human corneal barrier under dynamic flow blinking stimuli. This platform will open further possibilities in the field of ocular drug development and in future ocular mechanobiology studies.

## Conclusions

In this study, we established a biomimetic multi-corneal barriers-on-a-chip for testing the effect of shear stress forces mediated by eye blinking on the characteristics of the corneal epithelial barrier. The microfluidic device consisted of four chambers, and each chamber had an upper channel and lower channel separated by a clear PET porous membrane. The cell culturing compatibility of the microfluidic device was tested in HCE-T cells. This design allows the generation of the epithelial barrier characterized by both apical and proximal structures in the microfluidic device. The corneal epithelial barrier was formed after seven days of culturing in which ZO-1 expression was significantly upregulated. The permeability rate of fluorescein sodium in the microfluidic device was equivalent to that in the transwell system. The application of dynamic flow stimuli generated a shear stress of 0.6 dyn s cm^-2^. These forces did not affect cell adherence after 24 h. On the other hand, the dynamic flow stimuli increased the expression of CK-19 after 24 h. Moreover, t-SNE analysis indicated a clear separation between the cell population exposed to dynamic flow stimuli and that kept in static conditions. Furthermore, there was an increase in cell area, and the aspect ratio indicated a morphological change and presumably a phenotype alteration under the shear stress condition mediated by blinking recapitulation. Cells under dynamic flow stimuli presented flat and the high expression of the intermediate filaments of CK-19 assisted and supported the cell stretching process.

The multi-corneal barriers-on-a-chip provides a prove of the concept that the shear stress forces mediated by eye blinking do signal the superficial cells of the corneal barrier to alter their characteristics so they can cope with the repeatable mechanical stimuli during eye blinking. Therefore, dynamic flow stimuli must be considered in the development of future micro-physiological models of the human cornel barriers. In this context, our microfluidic device can provide a platform for future applications regarding ocular drug permeability, ocular drug toxicity, and in situ ocular mechanobiological studies.

## Supporting information

Supplemental informations

## Acknowledgments

Funding was generously provided by the Japan Society for the Promotion of Science (JSPS; 17H02083) and the Japan Agency for Medical Research and Development (AMED; 17937667). The WPI-iCeMS is supported by the World Premier International Research Centre Initiative (WPI), MEXT, Japan. The authors would like to thank Dr. Yoshikazu HIRAI for his kind support.

