## Supplemental informations for "Multi-corneal barriers-on-a-chip to recapitulate eye blinking shear stress forces"

**Supplementary Table. 1 The aspect ratios of HCE-T cells under static and flow dynamic conditions**. Data represent as mean ± SD of three independent experiments. *P*-Value was determined by the Nested T test.

| | **STATIC** | | | **DYNAMIC** | | | **P-VALUE** |
|---|---|---|---|---|---|---|---|
| **AREA** | A1 | A2 | A3 | A1 | A2 | A3 | <0.001 |
| **ASPECT RATIO** | 1.476±0.32 | 1.467±0.32 | 1.458±0.30 | 1.503±0.35 | 1.508±0.36 | 1.514±0.35 | |

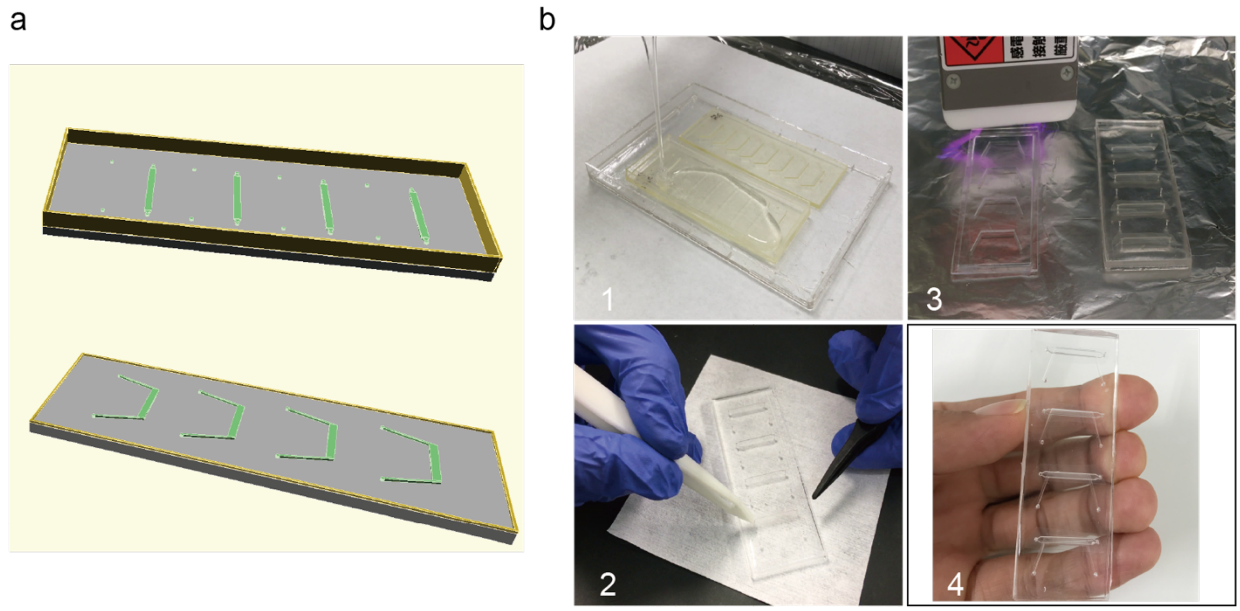


**Supplementary Fig. 1 The design and fabrication of the multi-well microfluidic device. a,** Design of the lower and upper channels with OpenSCAD. **b,** The process of fabrication; 1) The addition of PDMS and curing at 80 °C for overnight 2) The insertion of PET membranes in between the upper channels and lower channels 3) Bonding with corona treatment 4) The final structure of the multi-well microfluidic device**.**

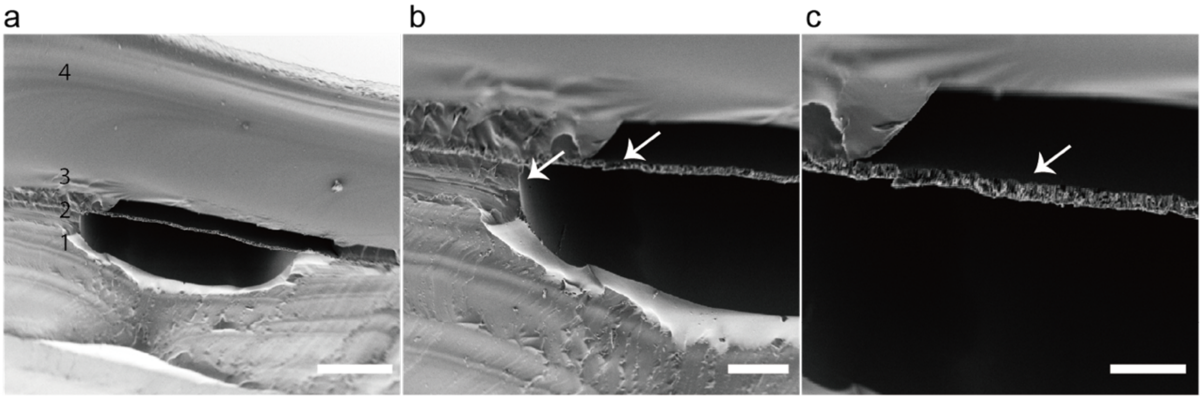


**Supplementary Fig. 2 SEM images of a cross-section of the microfluidic channel. a,** The structure of the microfluidic channel 1) Upper PDMS channel 2) PET porous membrane 3) Lower PDMS channel 4) Glass slide. Scale bar, 500 µm **b and c,** The intactness of the porous membrane after fabrication. Arrows indicates no gaps between upper and lower channels. Scale bar, 200 µm; 100 µm respectively.

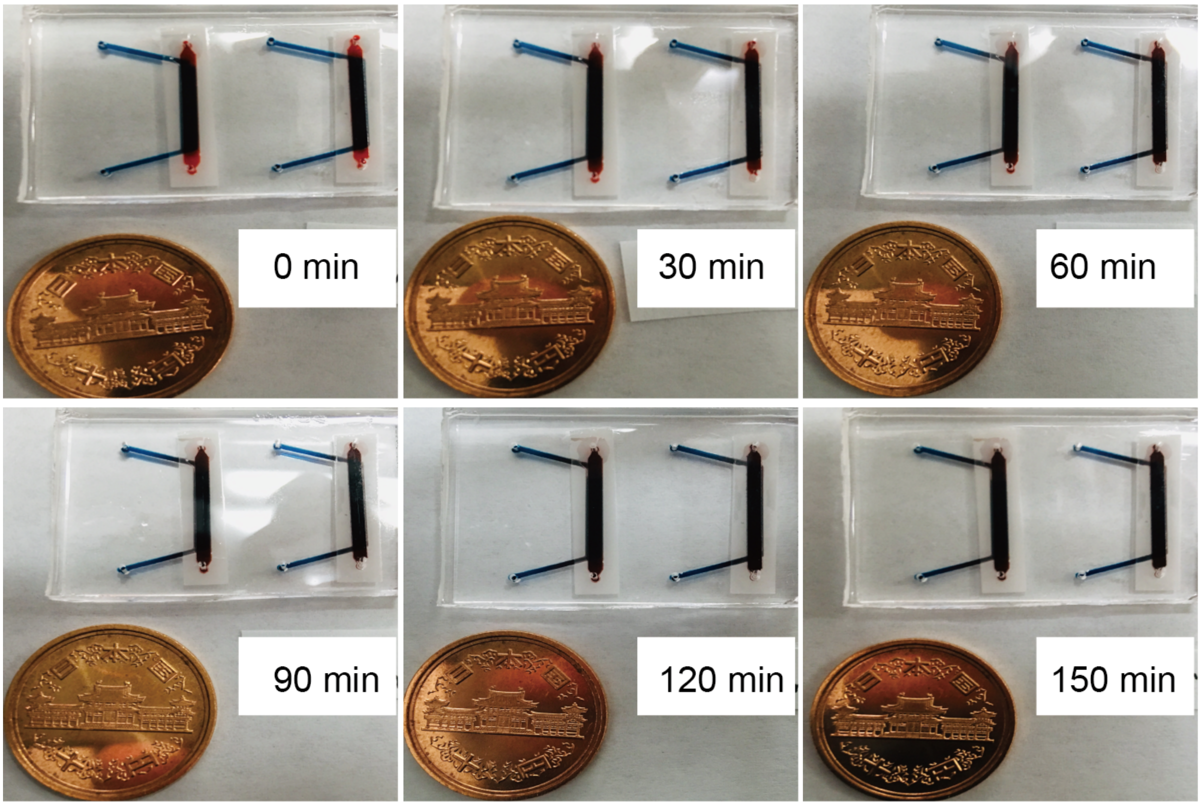


**Supplementary Fig. 3 Dye leakage and translocation in the multi-well microfluidic device.**

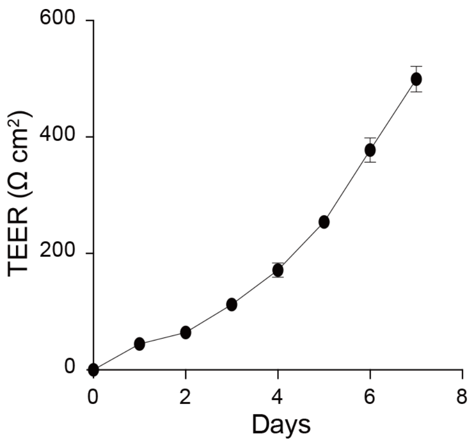


**Supplementary Fig. 4. TEER measurement of the corneal epithelial barrier in trans-well system.** HCE-T cells were seeded in trans-well in a density of 0.89 × $10^5$ cells $cm^{-2}$ for 7 days. Data represent as mean ± SD of eight independent experiments.

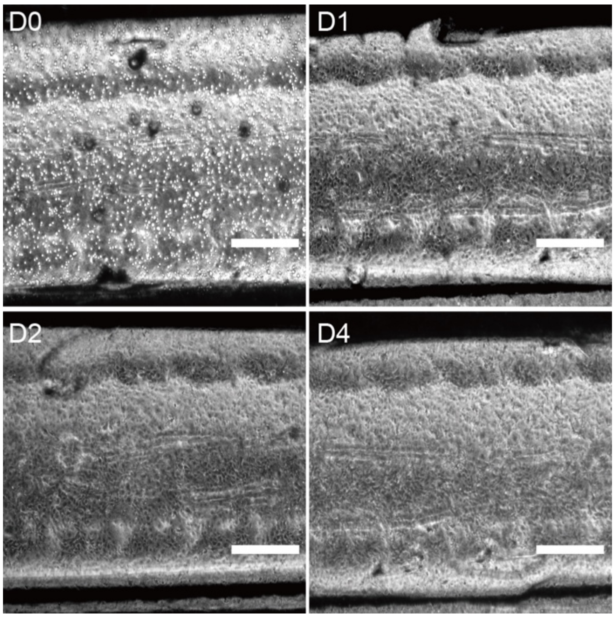


**Supplementary Fig. 5. Brightfield images of HCE-T cells post-seeding the multi-well microfluidic devices.** HCE-T cells were seeded in the microchannels in a density of $6 \times 10^3$ cells $cm^{-2}$. Scale bar, 500 µm.
